# Anterior inferotemporal cortex represents decision formation and termination without ramps

**DOI:** 10.64898/2025.12.24.696440

**Authors:** Gouki Okazawa, Roozbeh Kiani

## Abstract

Visually-guided decisions are often modeled as bounded accumulation of sensory evidence, with “ramping” activity in frontoparietal cortex taken as a canonical signature of integration. Here we show that anterior inferotemporal cortex (TEa) implements an alternative integration code during object-based decisions. In macaques performing stochastic face categorization, TEa neurons tracked momentary feature fluctuations while concurrently encoding an accumulated decision variable as a running average of task-relevant evidence. This running-mean code increased category signal-to-noise without ramps, supported flexible amplification of task-relevant features, and showed a sharp collapse in sensitivity to new evidence after commitment. These signatures emerged concurrently with, or slightly earlier than, those in simultaneously recorded LIP neurons, arguing against inheritance from feedback. A simple subunit model combining momentary evidence with a running-average decision variable accounts for TEa dynamics. Our results identify a previously unrecognized format for evidence integration and reposition TEa as a decision-aware hub within the ventral visual pathway.

## Introduction

Successful interactions with the visual world depend on two tightly coupled computations: forming representations from sensory input and transforming those representations into decisions that serve current goals. These computations are often studied in isolation and assigned to distinct cortical circuits. In classic random-dot direction discrimination, middle temporal (MT) neurons in the dorsal visual pathway encode momentary motion signals^1^, whereas frontoparietal regions such as the lateral intraparietal (LIP) neurons encode the integral of these signals over time to form decision variables that guide actions^2–6^. Traditionally, the same framework has been assumed to generalize to visual object recognition: the ventral visual pathway has been modeled primarily as a feedforward cascade that extracts features, sending representations of object identity to downstream frontoparietal circuits for decision-making and visually-guided behavior^7–9^. This division—sensory encoding in the ventral stream, decision formation elsewhere—has shaped both experiments and theory.

A central assumption in many accounts of decision formation is not only that evidence is integrated, but *how* that integration is expressed neurally. In frontoparietal circuits, integration is often associated with progressive, ramp-like changes in firing rates or population activity^2,4,10,11^. This “ramping” motif has become a de facto marker of accumulation. Yet integration need not imply ramps: mathematically, a decision variable can be represented in multiple formats that preserve or improve signal-to-noise ratio over time, including normalized or running estimates that can be updated online without maintaining a full history of samples^12–14^. Whether the brain exploits such alternative formats—and where—remains largely unresolved.

These questions are particularly salient for the inferotemporal (IT) cortex at the apex of the ventral pathway. Anterior IT (TEa) is richly interconnected with prefrontal, medial temporal, amygdalar, and striatal circuits^15–19^. Its pyramidal neurons have large somas, elaborate basal dendritic trees, high spine counts, and low myelination—properties more reminiscent of parietal and frontal association cortex than of earlier visual areas^20–23^. Consistent with this anatomy, IT responses exhibit rich and task-dependent dynamics: they evolve during viewing, converge onto category-specific patterns, and shift with task engagement and context^24–31^. Together, these observations raise the possibility that TEa does more than report object identity—it may participate in temporally extended computations that are required to decide.

Canonical models of perceptual decision making posit three core operations^4,12^. First, decisions form gradually as momentary sensory evidence is integrated over time windows extending hundreds of milliseconds to seconds^32,33^. Second, commitment to a choice occurs when the accumulated evidence reaches a threshold or bound, terminating deliberation^34–36^. Third, when stimuli vary along multiple dimensions, the system selectively integrates task-relevant features while discounting irrelevant ones^37–39^. These computations are well characterized in frontoparietal circuits^4,40^, but it remains unknown whether TEa expresses them—and, critically, in what representational format.

To directly probe how object representations contribute to decision formation, we used a face categorization task with stochastic fluctuations in facial features^10,41,42^. Unlike static object recognition paradigms or passive viewing tasks, our design allowed us to disentangle the encoding of momentary sensory evidence, its temporal integration into a decision variable, and decision termination (choice commitment) within the same trials and neuronal population. Our results reveal that TEa neurons (1) integrate evidence over time, (2) sharply reduce sensitivity to new input after commitment, and (3) flexibly amplify task-relevant visual features over irrelevant ones, dynamically adapting to changing task demands. Beyond these hallmark decision-making signatures, we identify a distinct format for the decision variable in TEa: instead of ramping, TEa encodes a running average of relevant evidence. This format yields a monotonic improvement in signal-to-noise ratio without classical ramps, and provides a compact code that can jointly support choice, confidence, and commitment^14^. These findings demonstrate that TEa, long viewed as a feedforward endpoint, contributes to decision computations in a representational format not previously emphasized in cortical decision-making circuits.

## Results

We recorded neural activity from TEa and LIP of two macaque monkeys as they performed a stochastic face categorization task^10^. On each trial, monkeys viewed a dynamic face stimulus and reported its category with a saccadic eye movement to one of two choice targets (Fig. 1A, B). Two variants of the task were tested in separate blocks: categorizing either face species (human versus monkey) or expression (happy versus sad). Each task used morphs between two prototype faces representing the two categories. The stimulus on each trial fluctuated stochastically every 106.7 ms around a nominal morph level (Fig. 1A inset; SD: 20% morph level; range ±100%, where +100% and −100% indicate the two prototypes). These within-trial fluctuations provided time-varying sensory evidence, allowing us to examine how the monkeys accumulated information to form a decision, and how IT neurons responded to fluctuating evidence. Stimulus durations varied across trials (227–1080 ms, mean 440 ms; truncated exponential), providing additional leverage to study decision dynamics across different integration windows^34^. Behavioral performance was consistent with bounded evidence accumulation, as confirmed by fits of a drift-diffusion model extended to accommodate multiple informative object features (Fig. 1C, D; see Methods; [10]).

**Figure 1:**
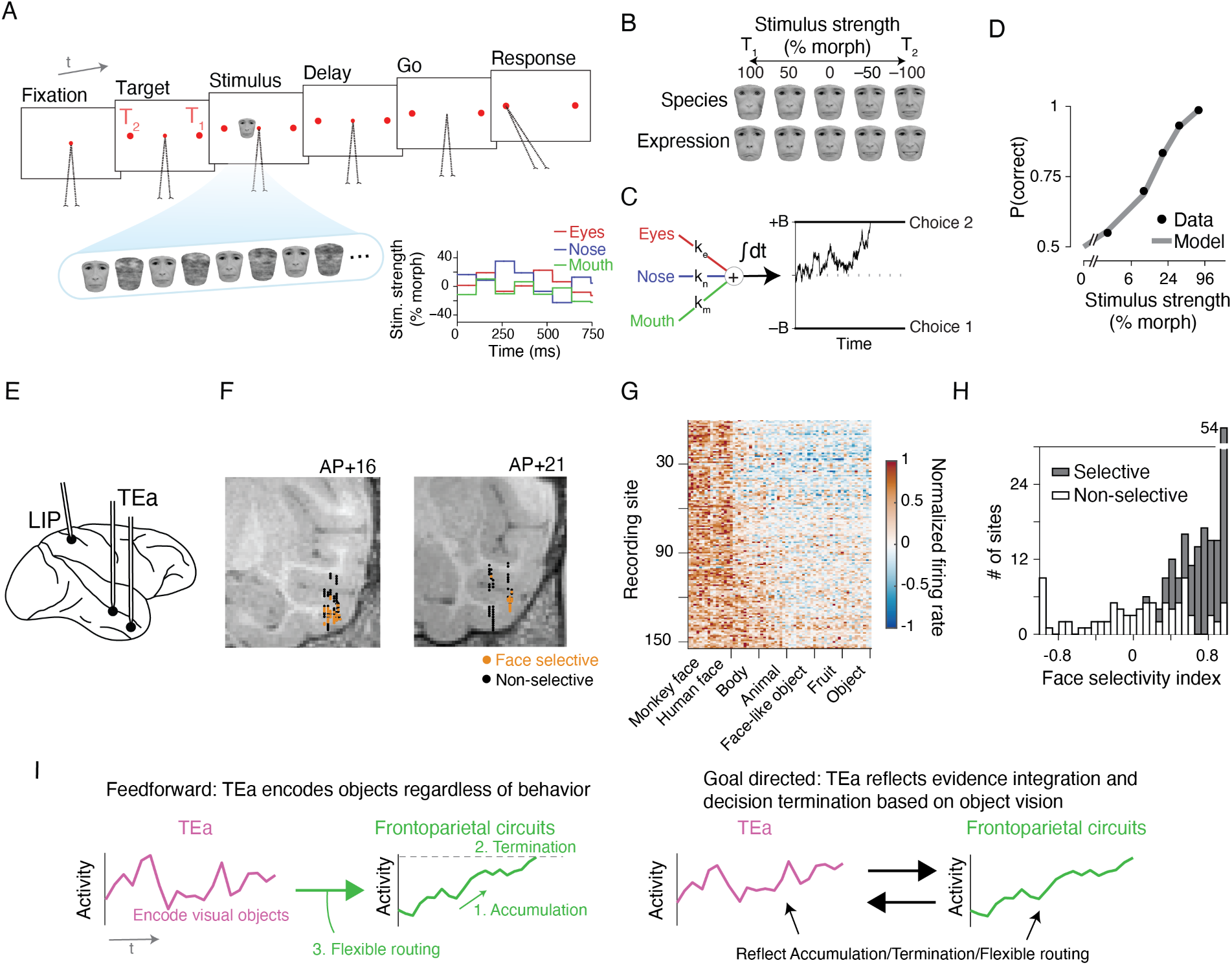
Face categorization task and neural recordings from IT and LIP. (**A**) Each trial began with fixation, followed by the appearance of two choice targets and a face stimulus. After the go cue (fixation point offset), the monkey reported the face category by making a saccade to one of the targets. During stimulus presentation, facial features stochastically varied every 106.7 ms around the trial’s nominal morph level (inset), providing dynamic evidence. The face fluctuations were interleaved with noise masks. The displayed human face images were obtained from Radboud database^43^ and used with permission for visualization purposes. (**B**) Monkeys discriminated either the species or expression of parametrically morphed face stimuli. (**C**) Behavior was well explained by an extended drift-diffusion model (DDM) that combined informative visual features into momentary evidence and integrated it over time^41,42^. (**D**) The monkey’s psychometric function and the DDM fit. (**E**) We recorded neuronal activity from the anterior inferotemporal (TEa) and lateral intraparietal (LIP) cortex during task performance. (**F**) In TEa, we targeted face-selective neural clusters, corresponding to face patches AL and AM, identified through electrophysiological mapping. (**G**) Face-selective sites (orange dots in F) showed stronger responses to faces than other object categories during passive viewing. (**H**) Large face-selectivity indices in face patches (see Methods), consistent with previous studies^44^. (**I**) Competing hypotheses about TEa computations: Under a feedforward framework (left), TEa is a sensory area that encodes sensory evidence, while downstream circuits accumulate evidence, terminate decisions, and flexibly route information. An alternative framework (right) emphasizes recurrent interactions, with decision-related signals distributed across both IT and frontoparietal regions.

During task performance, we simultaneously recorded from face-selective clusters and adjacent regions in TEa, as well as from LIP (Fig. 1E). To localize face-selective clusters in TEa, we first used a passive fixation task in separate blocks from the categorization task. In these blocks, we mapped the IT cortex with a recording grid and measured responses to face and non-face object images. In anterior IT of each monkey, we identified two face-selective clusters marked by consistently strong face-selective responses across contiguous recording sites (Fig. 1F). Based on stereotaxic coordinates (AP +16 and +21 in monkey 1; +20.5 and +22.5 in monkey 2) and anatomical landmarks (sulci and gyri on MR images), we identified these regions as face patches AL and AM^44^. For LIP, we targeted sites with robust delay-period selectivity as determined by a memory-guided saccade task performed in separate blocks from the categorization and passive fixation tasks^2,45^. Once we established target recording sites, we sampled them across consecutive sessions. Prior to each categorization block, we confirmed site selectivity in TEa and LIP using short blocks of passive fixation and memory-guided saccade tasks, respectively.

Figure 1G shows a heatmap of neural responses to face and non-face stimuli across sites, demonstrating strong face preferences in the targeted TEa clusters. This was quantified using a face-selectivity index (Fig. 1H; Eq. 1), consistent with previous reports^44^. Our multi-contact electrodes also sampled adjacent regions, which had weaker or no face preference (p > 0.05; Supplementary Fig. 1A) but were selective to other categories such as bodies. Across sessions, we isolated 367 units in TEa (187 from face-selective units and 180 from neighboring sites). Of these, 191 units (52%) showed selectivity for one of the face categories in the main task and were included in further analyses. Because neurons within and near the face patches exhibited similar task-related responses, we pooled these units in the main analyses. Results restricted to face-selective sites are shown in Supplementary Fig. 1B. For LIP, we recorded from 132 units, which showed clear selectivity for the saccade targets in the main task.

### TEa category encoding evolves over time

We first examined TEa neural responses to dynamic face stimuli in the categorization task. Single-unit and population-averaged peri-stimulus time histograms (PSTHs) revealed stereotyped dynamics: responses began ∼100 ms after stimulus onset, peaked at ∼200 ms, and gradually declined thereafter. Within this temporal profile, firing rates varied monotonically along the stimulus morph continuum (Fig. 2A, B). This selectivity emerged at response onset and persisted throughout stimulus presentation. Notably, for the preferred category of each neuron, responses to different morph levels gradually converged, reaching maximal overlap around 400 ms after stimulus onset (Fig. 2B; Supplementary Fig. 2A for individual monkeys). Such convergence—a hallmark of categorical encoding—is also characteristic of decision-related activity in frontoparietal circuits^35,36,46^. In contrast, no such convergence occurred for non-preferred categories, consistent with another well-established property of decision-making circuits and models: selected choices yield categorical representations, while unchosen alternatives retain graded encoding^35,47–50^. The convergence for the preferred category was evident across both short and long stimulus durations (Supplementary Fig. 3A), confirming that it was not an artifact of early truncation on short-duration trials.

**Figure 2:**
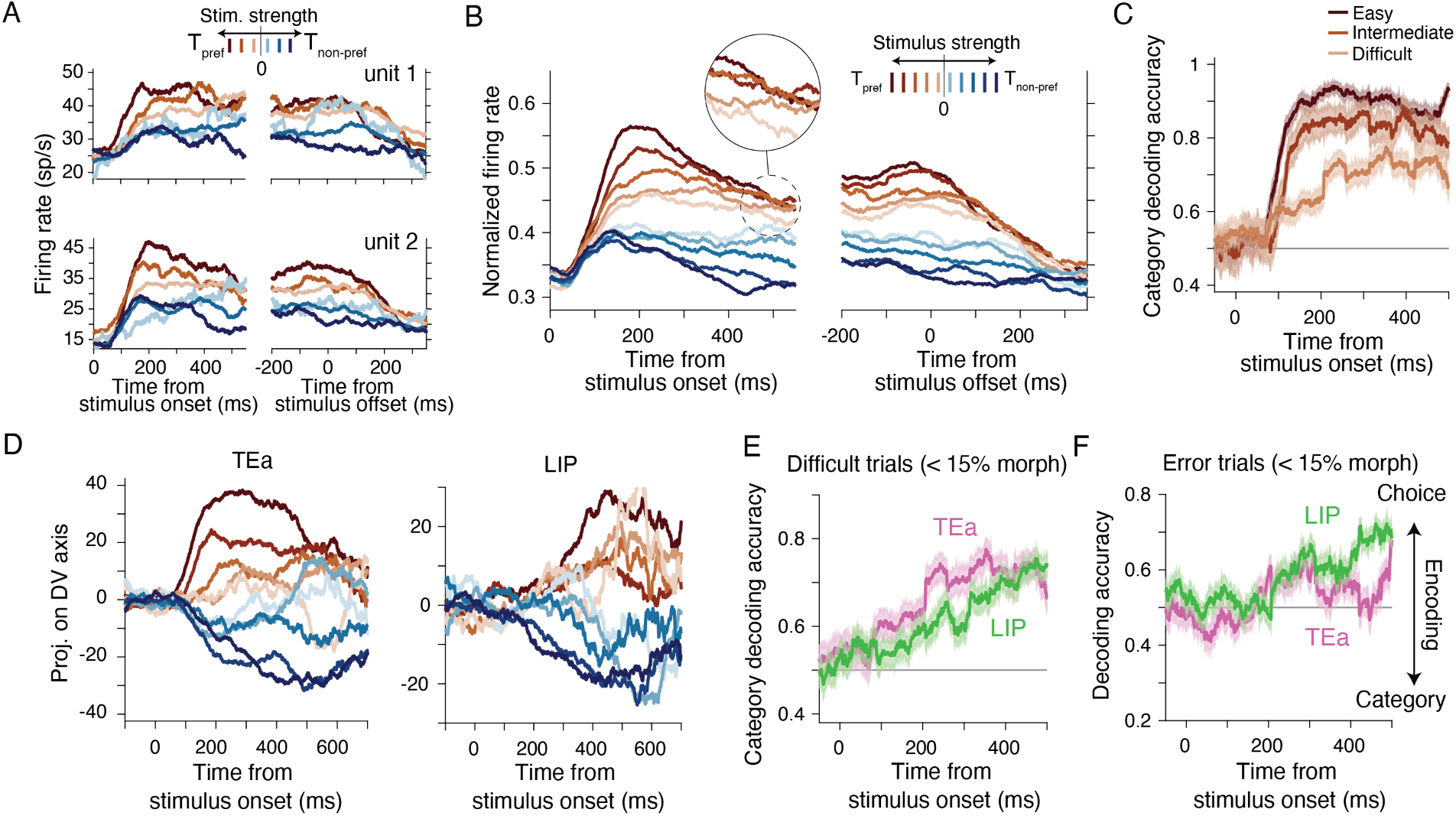
TEa activity reflects evidence accumulation. (**A**) Responses of example TEa neurons to different stimulus strengths during stimulus presentation. (**B**) Population-averaged responses of TEa neurons (n = 191) show monotonic tuning to stimulus strength. For the preferred category, responses to different stimulus strengths gradually converged after ~400 ms (inset). The plot includes correct trials only. (**C**) Category decoding accuracy from TEa population activity increased and saturated rapidly for easier stimuli, but ramped gradually for difficult stimuli. Shading indicates S.E.M. estimated by bootstrap. (**D**) TEa and LIP population activity projected on the axes encoding the decision variable, identified using canonical correlation analysis^10^. LIP: n = 132. (**E**) Category decoding accuracy ramped slightly earlier in TEa than LIP, challenging the hypothesis that IT decision signals arise from frontoparietal feedback. (**F**) To dissociate decision signals from sensory encoding, we analyzed error trials where stimulus category and monkey’s choice diverged. A classifier trained on correct trials preferentially decoded the monkey’s choice in error trials, with similar dynamics in TEa and LIP.

A hallmark of evidence accumulation is the gradual improvement in the signal-to-noise ratio of the decision variable. To assess whether TEa neurons exhibit this signature, or merely encode recent stimulus frames, we conducted a population-level classifier to decode category identity from single-trial activity patterns, using template-matching to population responses for each category (see Methods; [51, 52]). For easy stimuli, classification accuracy rose quickly and saturated just before the peak response, remaining high for the rest of the stimulus presentation (Fig. 2C; Supplementary Fig. 2B for individual monkeys). For more difficult stimuli (morph level< 20%), where accumulation of sensory evidence is most beneficial, classification accuracy increased gradually over time (Fig. 2C)—even as firing rates declined after ∼200 ms—suggesting the encoding of a latent decision variable that evolves over time. We return to the format of this code and its relationship to firing rate modulations in later sections. The ramping accuracy persisted significantly longer for more difficult stimuli than the easier ones (p < 0.001, bootstrap test based on ramp fits; see Methods).

We next asked whether this evidence accumulation signature in TEa could be inherited through feedback from downstream decision circuits. We used canonical correlation analysis to identify TEa and LIP decision-variable manifolds, comparing the evolution of neural population trajectories along those manifolds^10^. In LIP, population activity evolved along a curvilinear decision-variable manifold^10^, and began to separate different stimulus strengths ∼200 ms after stimulus onset (Fig. 2D)—well after TEa response onset and coinciding with TEa peak activity (Fig. 2B). In TEa, population activity also evolved along a curvilinear manifold (Supplementary Fig. 3B), but separation by morph level emerged ∼100 ms earlier than LIP (Fig. 2D). Consistently, the TEa population classifier accuracy began rising sooner than in LIP (Fig. 2E). This temporal lead was most pronounced for easy stimuli (p < 0.001) but remained significant for difficult ones as well (Fig. 2E; p = 0.02, bootstrap test based on fitting a ramp function; see Methods), challenging a simple feedback explanation for TEa decision signals.

Further, both TEa and LIP activity more closely reflected the monkey’s choices than the actual stimulus category. Using population response templates constructed from correct trials, we classified error trials based on similarity to each template. Classification accuracy above 0.5 indicates that neural responses aligned more with the animal’s choice than the physical stimulus category. Both TEa and LIP showed classification accuracies above 0.5 starting ∼200 ms after stimulus onset and remained choice predictive during stimulus presentation (Fig. 2F; p < 0.001, bootstrap test), reinforcing the view that TEa signals reflect an evolving internal decision variable, not merely the stimulus or top-down modulations.

Together, these results demonstrate that TEa population responses exhibit canonical signatures of evidence accumulation: increased encoding precision over time, response convergence for the chosen category, and choice-predictive dynamics that emerge in parallel with—and slightly ahead of—those in LIP. Even as firing rates decline, TEa representations become progressively more categorical, consistent with an active role in decision formation. In the following sections, we investigate how the stereotyped response dynamics of TEa neurons—an early peak in firing rate followed by a decline (Fig. 2A, B)—reflect evidence accumulation and termination.

### Decision termination in TEa is marked by reduced sensitivity to late evidence

The analyses above suggest that TEa does not operate as a passive sensory filter that merely encodes moment-to-moment visual input independent of ongoing decisions. To investigate the underlying computations, we examined how TEa neurons responded to within-trial fluctuations in sensory evidence. In our task, the morph level of the face stimulus varied every 106.7 ms around a fixed, trial-specific nominal mean, producing a sequence of discrete evidence “pulses” (Fig. 1A). To isolate the effect of each pulse, we subtracted the trial’s nominal mean morph level from individual pulses, yielding residual morph levels that were uncorrelated across time within a trial. We also computed residual spike counts by subtracting, at each time, the mean spike counts across trials with similar nominal morph levels. We then regressed these residual spike counts against the residual morph levels using a multivariate linear model (Fig. 3A; Eq. 6). The resulting regression weights are a measure of neural sensitivity, quantifying how strongly each pulse influences stimulus-dependent response dynamics.

**Figure 3:**
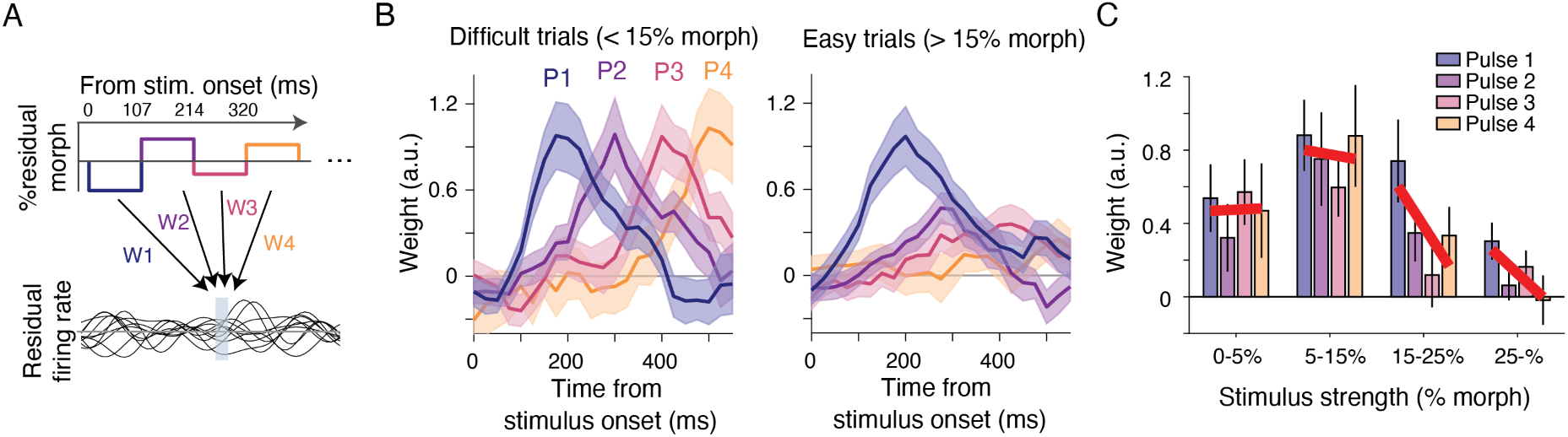
TEa activity reflects decision termination. (**A**) We leveraged frame-to-frame stimulus fluctuating (Fig. 1A, inset) to quantify neuronal sensitivity to momentary sensory evidence. Residual firing rates, obtained by subtracting time-dependent mean responses for each morph level, were regressed against residual morph fluctuations. (**B**) On difficult trials (left), TEa responses to each stimulus pulse remained robust and consistent over time, whereas on easy trials (right), responses were strong for the first pulse but diminished for subsequent pulses. Shading indicates S.E.M. across units. (**C**) Peak sensitivity for the first four stimulus pulses shown across difficulty levels. Thick red lines are linear fits to the four peak sensitivities. Sensitivity to stimulus fluctuations remained stable across pulses for difficult stimuli (<15% morph), but declined sharply for easy stimuli (>15%). Error bars indicate S.E.M. across units.

We found that the dynamics of neuronal sensitivity to stimulus pulses depended systematically on stimulus difficulty. On difficult trials (<15% morph), each stimulus pulse evoked a robust response, peaking ∼200 ms after pulse onset and decaying gradually over the next 200 ms (Fig. 3B left). In contrast, on easier trials (>15% morph), IT responses were strong for the first pulse but got substantially weaker for subsequent pulses (Fig. 3B right; Supplementary Fig. 2C for individual monkeys). To quantify this difference, we grouped trials into four stimulus difficulty levels and examined the peak regression weights for the first four stimulus pulses (Fig. 3C). For difficult stimuli, peak pulse sensitivities remained relatively constant across time. For easier stimuli, however, peak sensitivity declined monotonically across pulses, indicating a systematic reduction in the influence of later sensory evidence (p = 0.005, two-tailed *t*-test between the slopes of peak between easy and difficult trials).

Reduced sensitivity to late stimulus pulses on easier trials cannot be attributed to saturation of neural activity, as firing rates during the third and fourth pulses were well below peak firing rates observed earlier in the trial (Fig. 2A-B). Moreover, this pattern was specific to the preferred category, where categorical convergence of neural responses at decision termination is expected to reduce responses to later pulses, and was absent for the non-preferred category (Supplementary Fig. 4A), despite the lower firing rate bound at 0 spikes/s. These findings suggest that TEa ceases to encode new sensory evidence for the chosen category once sufficient information has been accumulated to commit to a decision.

Indeed, analysis of the monkeys’ behavior confirmed that late stimulus pulses influenced decisions on difficult trials more than easy trials (Supplementary Fig. 4B). We used psychophysical reverse correlation to quantify the impact of each pulse on choice likelihood^34,41,53,54^. On easy trials (>15% morph), reverse correlation dropped sharply after the first stimulus pulse and remained near zero from the third pulse onward, indicating minimal influence of later evidence. On difficult trials (<15% morph), reverse correlation declined gradually but remained significantly above zero, suggesting ongoing integration of sensory evidence. Notably, the rate of decline matched predictions from a bounded accumulation model as shown previously^41^: as the decision variable approached threshold, the marginal impact of later pulses diminished. The quantitative match between behavioral sensitivity and model predictions, and their similarity with neuronal responses support a unified account in which TEa neurons represent decision formation by integrating sensory evidence and ceasing to encode further input once a decision bound is reached.

### Mixed Encoding of momentary and accumulated evidence explains TEa response dynamics

Our results suggest that TEa responses reflect both the accumulation of sensory evidence and the termination of that process (Fig. 4A). However, TEa neurons do not show ramping activity during stimulus presentation (Fig. 2A, B). To reconcile this discrepancy, we investigated the representational format of TEa activity to discover how it might jointly encode momentary and accumulated evidence.

**Figure 4:**
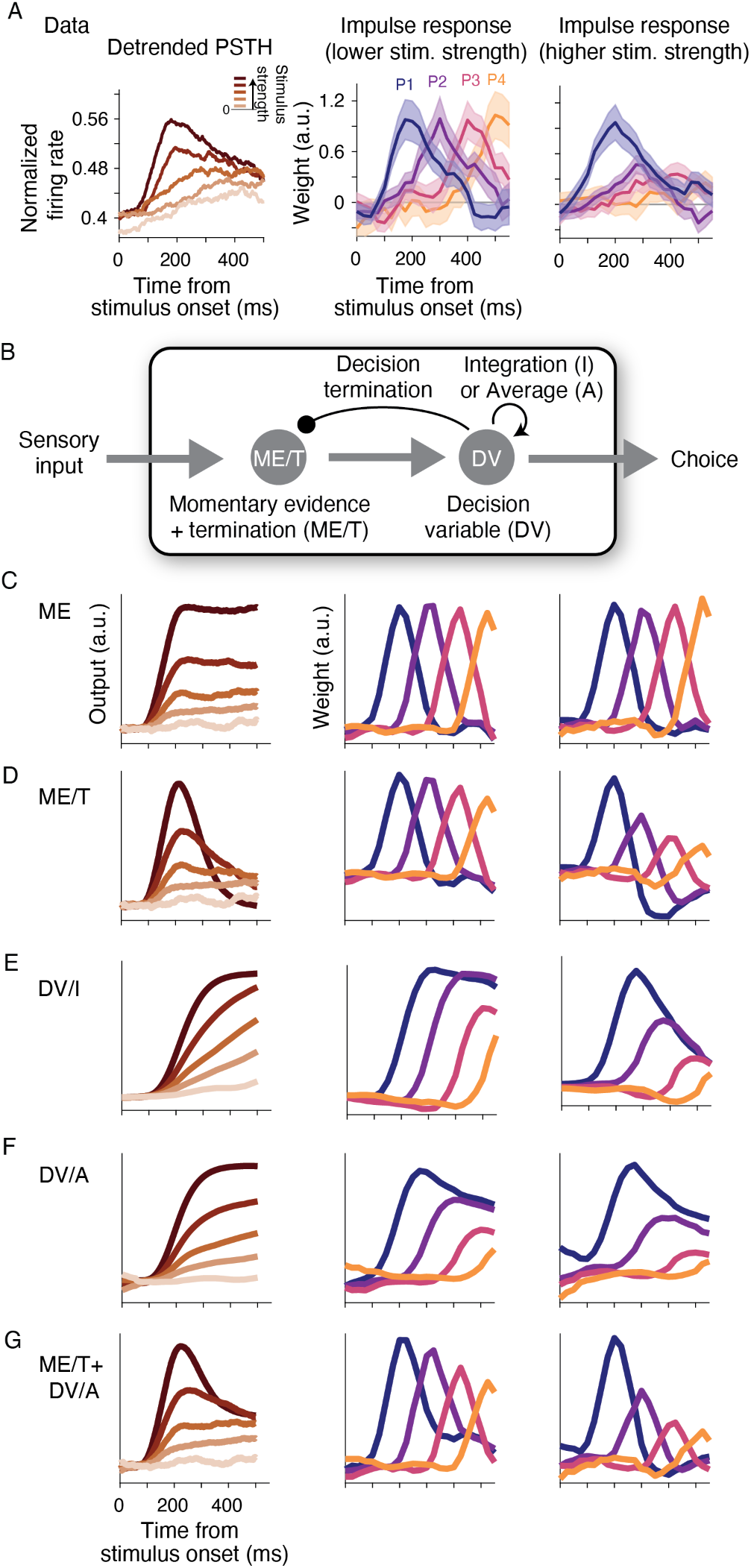
TEa responses are explained by joint encoding of momentary and accumulated evidence. (**A**) Summary of key features observed in the TEa population. Left: population-averaged PSTHs show gradual convergence. For visualization, PSTHs were detrended and plotted for the preferred category only. Middle and right: responses to individual morph fluctuations, replotted from Fig. 3B, reveal diminished sensitivity to late pulses in easy trials. (**B**) A simplified model consisting of two sub-units: one encoding the most recent visual input as momentary evidence (ME) and another accumulating task-relevant evidence into a decision variable (DV). (**C**) Without the termination mechanism, the ME sub-unit continues responding to stimulus fluctuations regardless of stimulus difficulty. (**D**) An alternative ME sub-unit with a termination mechanism that suppresses responses once the DV reaches a decision bound. This sub-unit has suppressed late responses but also exhibited a non-monotonic ordering of PSTHs across morph levels. (**E**) The DV/I sub-unit, which encodes the decision variable as an integral, increases its output over time (left) and maintains a lasting effect of early morph fluctuations (middle and right). (**F**) An alternative DV sub-unit, DV/A, that encodes the decision variable as a running average (i.e., the integral divided by elapsed time). (**G**) A hybrid model combining the ME/T and DV/A sub-units best captures the key features of TEa activity, including PSTH convergence and diminished sensitivity to late pulses, especially on easy trials.

We constructed a minimal model comprising two sub-units: one encoding momentary evidence (ME) and another integrating ME to create a decision variable (DV) (Fig. 4B). The ME sub-unit represented moment-to-moment stimulus fluctuations with a response profile that peaked 150 ms after each stimulus pulse (Gaussian temporal profile; SD, 20), with peak magnitudes proportional to the pulse strength. The output of the ME sub-unit, combined with additive Gaussian noise, was passed to the DV sub-unit, which accumulated task-relevant evidence until a decision bound was reached. We explored several variants of these sub-units to identify if they replicated the dynamics of TEa single-unit and population activity.

The base ME sub-unit responded to all stimulus pulses, regardless of their task relevance or when the decision-making process terminated. Consequently, its activity scaled monotonically with morph level and did not show the convergence of firing rates observed in TEa for the preferred category (Fig. 4C, left). Subjecting the activity of this ME sub-unit to our sensitivity analysis (Fig. 3) revealed uniform sensitivity across pulses, independent of difficulty or pulse timing (Fig. 4C, middle and right). Introducing a termination mechanism (ME/T), in which responses were suppressed after DV bound crossing, reduced sensitivity to later pulses, especially at high morph levels. However, this also produced an abrupt drop in activity following a peak at ∼200 ms (Fig. 4D, left) and yielded a reversal of sensitivity for the first two pulses of easy stimuli (Fig. 4D, right) This reversal arose because large early inputs drove the DV to its bound, suppressing the late phase of response to the same pulse (negative sensitivity).

We also tested two DV sub-unit variants. The first, DV/I, directly integrated task-relevant momentary evidence, representing the decision variable in a format similar to standard decision-making models (e.g., DDM; [12, 55, 56]) (Fig. 4E, left). The sensitivity of DV/I sub-unit declined for later pulses across the trial, most saliently for easy stimuli. Further, because the DV/I sub-unit maintained an integral of evidence, its sensitivity to earlier pulses persisted throughout stimulus presentation, especially for the weaker stimuli where multiple pulses were required to drive the DV to its bound (Fig. 4E, middle and right).

The second variant of the DV sub-unit, DV/A (Fig. 4F), did not represent the integral directly. Rather, it estimated the overall stimulus morph level, computing the running average of task-relevant ME sub-unit outputs. Since the average equals the integral divided by elapsed time, the DV/A sub-unit indirectly represented the integration process. The outputs of both the DV/I and DV/A sub-units are equally suitable for identification and categorization tasks, as both reflect the gradual improvement of signal-to-noise ratio through evidence integration across pulses. Also, both show reduced influence of the late pulses, especially at high morph levels. However, the DV/I sub-unit output matches standard bounded evidence accumulation models and is suitable for decision termination with a fixed bound, whereas the DV/A sub-unit output converges to the mean stimulus morph level and requires a time-varying bound to implement a decision threshold^14^.

None of the sub-units alone captured the full pattern of TEa response properties (Fig. 4C-F). However, their limitations were complementary, suggesting that a mixed coding model may better explain the data. Specifically, a hybrid model combining ME/T and DV/A matched single unit and population response dynamics very well (Fig. 4G). Models with the base ME sub-unit consistently overestimated sensitivity to late pulses, while models with the DV/I sub-unit underestimated the convergence of responses across morph levels of the preferred category (Supplementary Fig. 5). By contrast, the ME/T+DV/A model reproduced key patterns of TEa responses, including a peak at ∼200 ms after stimulus onset, diminished sensitivity to late pulses at high morph levels, and convergence of firing rates across morph levels later in the trial (Fig. 4G). Although this hybrid model focuses on responses to the preferred stimulus category (Fig. 4A) and requires additional sub-units to account for asymmetries between preferred and non-preferred categories (Fig. 2B, Supplementary Fig. 6), it demonstrates the necessity of mixed coding schemes to explain the decision-related dynamics observed in TEa. Overall, TEa activity represents a mixture of the momentary evidence from individual pulses, a decision variable formatted as a running average across pulses, and the termination of evidence accumulation.

While the DV signal in TEa does not ramp, it nevertheless carries functionally equivalent information for categorization tasks. Our hybrid models suggest that TEa encodes both “what is” and “what has been,” and ceases updating once a sufficient level of evidence—*quantum satis*—is accumulated.

### Decision-related TEa activity is context dependent

While IT neurons are known to encode multiple stimulus features^57–59^, our hypothesis that they encode the decision variable predicts that they should preferentially represent the feature relevant to the current task. To test this context dependence, we analyzed TEa neurons recorded while monkeys performed the species and expression categorization tasks within the same session. The stimuli were sampled from a two-dimensional (2D) face space defined by orthogonal morph axes for species and expression (Fig. 5A). Critically, stimulus fluctuations occurred along both axes in each task, allowing us to compare TEa responses to identical visual inputs under different task rules.

**Figure 5:**
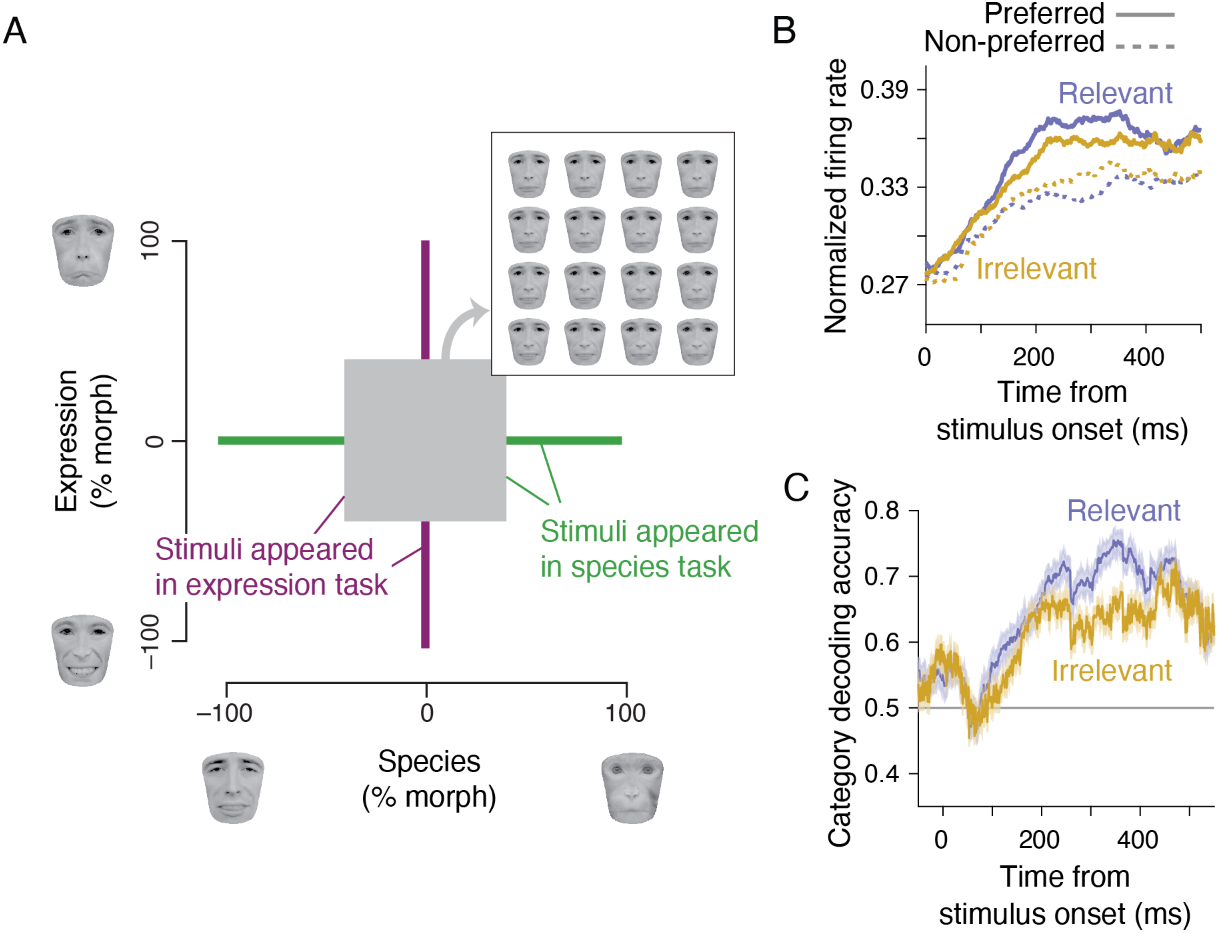
TEa activity is context dependent. (**A**) Monkeys performed species and expression categorization tasks using overlapping stimuli (central gray square and inset). This design enabled us to test whether TEa responses to identical visual stimuli differed across task contexts. The displayed human face images were obtained from Radboud database^43^ and used with permission for visualization purposes. (**B**) The difference in neural responses between preferred and non-preferred categories was greater for task-relevant than task-irrelevant stimulus dimensions. (**C**) Population-level classification accuracy was higher for task-relevant categories, even when using the same units and the same stimuli. Shading indicates S.E.M. estimated by bootstrap. See Supplementary Fig. 7 for individual monkeys.

We found that TEa neurons preferentially encoded stimulus fluctuations along the task-relevant axis, beginning 100-200 ms after stimulus onset. Population-averaged PSTHs of the same stimuli in the two tasks revealed greater separation between preferred and non-preferred categories along the task-relevant axis than the irrelevant axis (Fig. 5B; t = 2.67, p = 0.0087, two-tailed *t*-test; Supplementary Fig. 7 for individual monkeys). To quantify the functional impact of this preferential code, we trained population-level classifiers. Classification accuracy improved more strongly for task-relevant categories than for task-irrelevant ones (Fig. 5C; p < 0.001, bootstrap test).

These results indicate that TEa encoding is not fixed but flexibly modulated by task demands. Rather than passively reflecting visual features, TEa dynamically prioritizes task-relevant information, consistent with an active role in transforming visual inputs into decision variables. However, this preference is not absolute; irrelevant stimulus fluctuations continued to modulate TEa activity, albeit more weakly than the relevant fluctuations (Fig. 5B-C). This pattern of concurrent but differential encoding of task-relevant and irrelevant inputs aligns with our hybrid model of TEa neurons (Fig. 4G): the ME sub-unit encodes all stimulus fluctuations while the DV/A sub-unit encodes the integral of task relevant information as a running average.

## Discussion

We identified a previously unrecognized role for the anterior inferotemporal cortex (TEa) in perceptual decision making, and, critically, a previously underappreciated format for neural evidence integration. Leveraging a stochastic face categorization task, we show that TEa encodes momentary fluctuations in visual evidence while simultaneously representing an accumulated decision variable and the termination of accumulation. These decision-related signatures—temporal integration of task-relevant evidence, commitment-associated suppression of sensitivity to late evidence, and preferential encoding of task-relevant features—are typically attributed to frontoparietal decision circuits. Our results therefore revise a common view of IT as a static perceptual buffer and instead place TEa as an active computational node that shapes visual representations in the service of behavior.

A key advance is that TEa represents accumulated evidence in a non-ramping format. In frontoparietal regions, evidence accumulation is commonly linked to ramp-like changes in firing rates or population activity^4,11,12^. In contrast, TEa neurons do not exhibit classical ramps at the single-cell level. Yet, TEa nonetheless carries a decision variable whose signal-to-noise improves over time, both in individual neurons and across the population. Our analyses and modeling indicate that TEa encodes this variable as a running average of momentary evidence. This format preserves the functional consequences of integration while producing stereotyped temporal profiles—early peaks followed by declines—rather than sustained ramps.

The running-average code offers a principled computational account for how TEa can support decision formation at the apex of the ventral stream. First, the running average is mathematically tied to the integrated evidence: it equals the integral divided by elapsed time (or pulse count in our task), and can be updated recursively without storing the full stimulus history^13,14^. This makes it an efficient online representation of accumulated evidence. Second, the code naturally supports a monotonic improvement in signal-to-noise during stimulus viewing, consistent with our decoding results, without requiring monotonic firing-rate ramps. Third, the running average provides a compact basis for computing decision confidence and commitment, and is naturally compatible with decision termination via a collapsing bound^14^. In this view, TEa provides a decision variable that is well matched to object categorization: a normalized estimate of category evidence that remains directly interpretable as stimulus identity while still supporting downstream decision computations.

Our task design was critical for identifying this format. Most object recognition paradigms use static stimuli, limiting the ability to separate momentary evidence from accumulated evidence and from commitment-related changes in sensitivity. By introducing stochastic fluctuations in diagnostic and non-diagnostic facial features, we could measure (i) sensitivity to momentary evidence at each time point, (ii) the gradual improvement in category coding expected from accumulation, and (iii) the collapse of sensitivity to late evidence after commitment. This combination—together with a subunit model—allowed us to distinguish a running-average decision variable from alternative accounts based on pure sensory encoding or integral-like accumulation of evidence.

The presence of decision-related computations in TEa aligns with its anatomical position and cellular properties. TEa sits at a strategic intersection of perceptual, mnemonic, emotional, and executive systems, receiving bottom-up input from ventral visual stream while maintaining reciprocal connectivity with medial temporal lobe structures (e.g., perirhinal cortex, entorhinal cortex, hippocampus) and prefrontal regions including ventrolateral and orbitofrontal cortex^15,16,19^. This convergence allows TEa to access both immediate sensory information and context-dependent modulatory signals related to memory, value, and goals. Moreover, TEa shares several architectonic features with frontoparietal decision-making circuits, including large somas, lower cell densities, elaborate basal dendritic trees, and high spine densities^20,21,23^. These specializations likely support recurrent processing and integration of distributed signals. Our results provide functional support for this view: the activity dynamics of TEa neurons exhibit temporally extended computations that go beyond momentary sensory encoding.

Although TEa encodes accumulated evidence, the mechanism that terminates the decision-making process may not originate locally. Recent studies point to the superior colliculus (SC) as a likely source of termination signals^36^. The SC also has privileged access to rapid visual signals that support face-selective responses^60^. This raises the possibility that TEa constructs the decision variable in a running-average format, while termination signals—reflecting threshold crossings, urgency, or action gating—are inherited from the SC. Such an architecture would explain the observed reduction in TEa sensitivity to late evidence and would align with the known role of SC in triggering saccades and decision commitment^36,61–63^.

Finally, our data suggest that decision variables may be reformatted across circuits for different computational demands. TEa activity reflects integration of evidence during stimulus viewing, but does not maintain this information across the post-stimulus delay^64^ (Supplementary Fig. 3C). In contrast, LIP neurons maintain decision-related signals through the delay, consistent with their role in motor planning and working memory. Given that LIP activity trails TEa and adopts a different representational format (ramping), a plausible working model is that TEa computes an early DV as a running average suited for object inference, which is then relayed and transformed into an integral-like code in frontoparietal circuits to support action-related computations and persistent choice representations^40,65^. A similar transformation may occur in the SC, which reflects both the DV and the commitment to motor responses^36,66^. This perspective positions TEa as a key node in transforming visual information into goal-directed behavior, and highlights that the neural “signature” of integration can vary systematically with circuit position and computational role.

## Competing interests

The authors declare no competing interests.

## Data availability

All data will be made publicly available upon publication.

## Code availability

Analysis code will be made publicly available on GitHub upon publication.

## Acknowledgments

This work was supported by the Simons Collaboration on the Global Brain, Pew Innovation Fund, and National Institute of Mental Health (R01MH127375, R01MH109180, and R01MH141929). G.O. was supported by postdoctoral fellowships from the Charles H. Revson Foundation and the Japan Society for the Promotion of Science.

## Methods

### Subjects and apparatus

We recorded neural activity from the anterior inferior temporal cortex (TEa) and the lateral intraparietal area (LIP) in two adult macaque monkeys (*Macaca mulatta*) as they performed face categorization tasks. All experimental procedures were approved by the Institutional Animal Care and Use Committee at New York University and conformed to the National Institutes of Health *Guide for the Care and Use of Laboratory Animals*.

Monkeys were seated in a semi-dark room with their heads stabilized using a surgically implanted head post, facing a cathode ray tube monitor (frame rate, 75 Hz). Stimulus presentation was controlled with Psychophysics Toolbox^67^ and Matlab. Eye position was monitored at 1 kHz using a high-speed infrared eye-tracking system (Eyelink, SR-Research, Ontario).

### Stochastic face categorization task

We trained monkeys to classify faces into two categories, defined by prototype faces. Each trial began when the monkey voluntarily fixated on a small point at the center of the screen (fixation point; diameter, 0.3°). Shortly afterward, two red choice targets appeared on opposite sides of the screen, equidistant from the fixation point (diameter, 0.5°; eccentricity 5°-12°). After a variable delay (range, 250–500 ms, truncated exponential distribution), a face stimulus appeared 1.6°–2.5° contralateral to the recording site. Stimulus size varied randomly across trials by one octave to discourage reliance on local features (range of stimulus width, 2.9°-5.8°; mean, 4.2°).

The stimulus was presented for a variable duration (range, 227–1080 ms; mean, 440 ms; truncated exponential), followed by a delay period (range, 300–900 ms; mean, 600 ms; truncated exponential). After the delay, the fixation point disappeared (Go cue), instructing the monkey to report the face category with a saccadic eye movement to one of the targets. Correct responses were rewarded with juice. To control task difficulty, we created a morph continuum between the two prototypes (see below). When the nominal morph level of the stimulus was halfway between the two prototypes, the monkey was rewarded randomly, irrespective of the choice.

Monkeys were trained to categorize face stimuli either by species (human vs. monkey) or expression (happy vs. sad) (Fig. 1B). The same set of stimuli was used for both tasks. We use the terms “happy” and “sad” descriptively, without implying that monkeys interpret expressions as humans do. The two tasks build on the monkey’s innate ability to discriminate between facial identities and gestures. Monkeys performed these tasks in blocks that switched either within or across sessions. To cue the task context at the start of each block, we presented stimuli near category prototypes. After a few trials, we introduced the full range of morph levels. In both categorization tasks, monkeys easily generalized across facial identities. We therefore created multiple stimulus sets from different human and monkey faces. Each set supported both categorization tasks (see below). We alternated between stimulus sets across sessions.

Each stimulus set formed a two-dimensional (2D) “face space,” defined by orthogonal morph axes for species and expression (Fig. 5A). To generate this space, we selected three “seed” faces: photographs of happy and sad expressions of a human face and neutral expression of a monkey face. The seed human faces were obtained from the MacBrain Face Stimulus Set^68^ (http://www.macbrain.org/resources.htm) and the monkey faces from the PrimFace database (http://visiome.neuroinf.jp/primface). We manually labeled 96–118 anchor points on each seed face and used a custom algorithm to morph the three seed faces by computing a linear weighted sum of the positions of the anchor points and textures inside the tessellated triangles defined by the anchor points^42^. Our algorithm allowed independent morphing of different stimulus regions. Further, because our morphing changed both the geometry and content of the facial regions, the resulting faces were perceptually seamless (Figs. 1 and 4).

Natural faces are complex and high-dimensional stimuli that challenge the investigation of the decision-making process. To control stimulus complexity, we limited informative features to three key regions: eyes, nose, and mouth, reducing the effective stimulus dimensionality to three. All other stimulus regions were fixed at the midpoint of the morph space and were therefore uninformative. Each informative feature varied within a 2D “feature space,” defined by the three seed faces. Morphed features were generated as weighted combinations of the three seed features. For species morphing, the weights for the happy (W*_h_*), sad (W*_s_*), and monkey (W*_m_*) seeds varied from [0.5, 0.5, 0] to [0, 0, 1], corresponding to morph levels −100% and +100% (species prototypes). For expression morphing, the weights ranged from [0.75, −0.25, 0.5] to [-0.25, 0.75, 0.5] (expression prototypes). Negative weights indicate linear extrapolation beyond the seed faces. We verified that for the weights used in our stimulus sets, all morphed features and resulting faces looked naturalistic and did not show noticeable aliasing or other abnormalities. In the 2D space, the species morph level of a feature was defined by (W*_m_* − W*_h_* − W*_s_*) × 100%, and its expression morph level was defined by (W*_s_* − W*_h_*) × 100%. The face space of Fig. 5A represents a 2D slice through the full 6D stimulus space, where the morph levels of three informative features are identical within individual stimuli but different across the stimuli.

On each trial, we selected a nominal morph level along the relevant task axis (species axis for species categorization or expression axis for expression categorization), determining both the target response and the mean morph level of the three features. Nominal morph levels ranged from –96% to +96% and were logarithmically spaced to approximate perceptually equi-distant stimuli. The exact set of nominal morph levels was adjusted for each stimulus set to keep the monkey’s overall accuracy constant across stimulus sets and tasks. To investigate how monkeys weighted sensory evidence conferred by the three informative features, we introduced stochastic fluctuations in the three features every 106.7 ms during stimulus presentation^41,42^. For all nominal morph levels between −12% and +12%, each feature fluctuated independently along both task-relevant and irrelevant axes according to a 2D Gaussian distribution (SD=20% morph level). Because of the logarithmic spacing of nominal morph levels, a large number of trials (42.4%) fell within this target region. For the remainder of the trials with stronger nominal morph levels (>12%), fluctuations were restricted to the relevant axis to prevent context confusion. Samples outside the prototype range [–100% +100%] were replaced with new samples inside the range (5.2% of cases).

We used a masking procedure to keep changes of features in a trial subliminal^41^. The masks were created by phase randomization of faces^69^ and were interleaved with the stimuli. Each 106.7 ms sample-mask cycle began with a face stimulus shown for one monitor frame, followed by fading to a mask. In the fading period, the mask and the stimulus were linearly combined, pixel-by-pixel, according to a half-cosine weighting function, such that in the last frame, the weight of the mask was one and the weight of the face stimulus was zero. In the next cycle, a new face stimulus with slightly altered informative features (Gaussian sampling) was shown, followed by fading with another mask, and so on. Before the first sample-mask cycle in the trial, we showed a mask frame to ensure all stimulus samples were preceded and followed by masks. The masking procedure was quite effective. In pilot human experiments, subjects could not detect feature changes, even though these changes influenced their choices^42^. The stimulus in each trial looked like a face appearing and disappearing behind cloud-like patterns. Each trial included 2-10 stimulus-mask cycles (mean, 4; truncated geometric distribution), yielding stimulus durations of 227–1080 ms (including the initial mask). An example video showing an image sequence used in the experiment can be found in Okazawa et al. (2021b).

Two monkeys performed the task across 80 recording sessions (monkey 1, 41 sessions; monkey 2, 39 sessions), which amounted to a total of 61,430 trials (monkey 1, 18,185 and 18,171 trials; monkey 2, 13,212 and 11,862 trials for species and expression categorization tasks, respectively).

### Neural recording

We recorded single and multi-unit activity from the inferior temporal cortex (IT) and the ventral division of the lateral intra-parietal cortex (LIPv) in the same two monkeys. In IT, we targeted face-selective clusters AL and AM^44^ and their neighboring regions (Fig. 1). In LIP, we focused on a subregion showing spatially selective persistent activity during a memory-guided saccade task (see below). Recordings were performed through plastic chambers (Crist Instruments, Damascus, MD) implanted on the skull, and a 1-mm spaced grid was placed inside the chambers for precise electrode targeting. We used either single tungsten microelectrodes (FHC, Bowdoin, ME) or 16-channel linear array probes (V-Probe; Plexon, Dallas, TX). Action potential waveforms were sorted offline (Offline Sorter, Plexon). We included both well-isolated single units and multi-units in the analyses.

Face-selective IT clusters were identified via extensive mapping of neural selectivities using single electrodes advanced through a grid with 1 mm spacing. While monkeys passively fixated, we presented either 66 or 96 colorful photographs of faces, bodies, animals, fruits, artificial objects, and face-like round objects (e.g., clocks), each shown for 106.7 ms against a gray background with 240 ms inter-stimulus interval. Firing rates were measured in a window from 70 ms after stimulus onset to 110 ms after stimulus offset^70^. Recorded sites were classified as face selective if their responses to face stimuli were significantly greater than non-face stimuli (*t*-test, p < 0.05; orange points in Fig. 1F). We found one face-selective cluster at the lateral convexity near the ventral lip of the superior temporal sulcus (putative face patch AL; monkey 1, AP +16; monkey 2, AP +20.5), and another more anterior along the lateral surface (putative face patch AM; monkey 1, AP +21; monkey 2, AP +22.5). To compare our recordings with previous studies^44^, we computed a face-selectivity index (FSI):

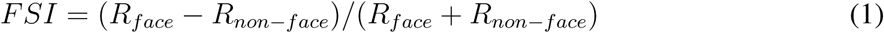

where R*_face_* and R*_non−face_* are the average firing rates to face and non-face images, respectively. After identification of these face patches, we repeatedly targeted them across sessions. At the beginning of each session, we confirmed electrode positions using a short fixation block prior to the categorization blocks.

For most analyses, we focused on the TEa neurons that showed significant category selectivity for the stimulus sets used in our categorization tasks. We measured firing rates during the stimulus period (from 80 ms after stimulus onset to 80 ms after stimulus offset) and identified task-selective units via a *t*-test comparing responses to the two face categories (p < 0.05). Of 367 total units (monkey 1, 229; monkey 2, 138), 191 (52%) met this criterion (monkey 1, 127; monkey 2, 64), including 95 units recorded within the face-selective clusters and 96 from the adjacent regions. Because results from units within the face patches were qualitatively similar to those in the adjacent regions (Supplementary Fig. 1B), we included all task-selective units in the main analyses.

For the context-dependence analyses in Fig. 5, we focused on units recorded in both tasks within the same sessions, selecting those that exhibited stimulus selectivity in either task and had an average firing rate greater than 5 spike/s during stimulus presentation (monkey 1, 61). These criteria prevented biasing unit selection toward those showing category selectivity along the task-relevant axis. For individual monkey analyses in Supplementary Fig. 7, we used a more inclusive criterion, selecting all units recorded in either task with an average firing rate above 5 spike/s during stimulus presentation (monkey 1, 153; monkey 2, 116).

LIP units were selected for analysis if they exhibited spatially-selective persistent activity during the delay period of a memory-guided saccade task^71^. While the monkey fixated centrally, a target briefly appeared at one of several locations, followed by a ∼1000 ms delay. At the end of the delay period, the fixation point turned off (Go cue), instructing the monkey to make a saccade to the remembered target location. The response field (RF) of each unit was identified as the target location eliciting the strongest delay-period activity. In the categorization task, one choice target (T_in_) was placed inside the RF and the other (T_out_) on the opposite side of the screen. We recorded 132 LIP units across the two monkeys (monkey 1, 62 units; monkey 2, 70 units).

### Data analysis

#### Behavioral data analysis

Psychometric functions (Fig. 1D) were calculated as the probability of making correct choices as a function of stimulus strength. Following our previous work^10^, we defined stimulus strength as the average morph level across stimulus-mask cycles and facial features in each trial. Trials were then grouped into five bins: 0–5%, 5–15%, 15–25%, 25–50%, 50–100% morph.

To test if the behavioral performance was consistent with evidence accumulation, we fit a drift diffusion model (DDM) to the data^10,41,42^. In the DDM, noisy momentary sensory evidence is accumulated over time to form a decision variable (DV). At each time point t, momentary evidence is drawn from a Gaussian distribution with standard deviation σ = 1 and mean µ(t). This mean was a weighted sum of the morph levels of the three informative facial features:

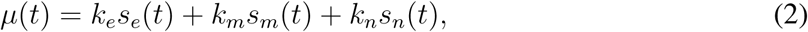

where s*_e_*(t), s*_m_*(t), and s*_n_*(t) are morph levels of the eyes, mouth, and nose along the task-relevant axis, and k*_e_*, k*_n_*, k*_m_* are weights reflecting sensitivity to each feature. Accumulation of momentary evidence continued until the DV reached either an upper (+B) or a lower bound (−B) or until stimulus offset. If a bound was reached, the corresponding choice was made; otherwise, the sign of the DV at stimulus offset determined the choice. We fit the DDM to each monkey’s choices using maximum-likelihood estimation. Full modeling details are described elsewhere^41^.

To quantify how monkeys used sensory evidence over time, we performed psychophysical reverse correlations^41^. Psychophysical kernels K(t) were calculated as the difference in average morph level fluctuations conditional on choice:

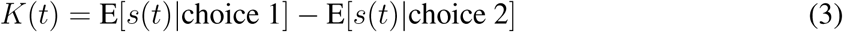

where s(t) represents the morph level at time t. For trials with nonzero average morph levels, the mean morph level was subtracted to focus on residual fluctuations^41^. To assess dependence on stimulus strength, we compared kernels for trials with low (0-15%) and high (15-40%) average morph levels (Supplementary Fig. 4B). The range for the high morph group was chosen to ensure that calculation of residuals was not affected by the bounded range of morph line. The morph level fluctuations of the three features were averaged using the weights derived from the DDM fit (eq. 2).

#### Generation of PSTHs

Peri-stimulus time histograms (PSTHs) were computed by averaging spike trains across trials, grouped by stimulus strength. Trials were sorted into five bins based on the average morph level across stimulus-mask cycles, as in the behavioral data analysis. PSTHs were aligned to stimulus onset or offset and smoothed by convolution with a 100-ms boxcar filter.

For population-averaged PSTHs, we defined the preferred task-relevant category of each unit as the category eliciting a higher mean activity during stimulus viewing (from 80 ms after stimulus onset to 80 ms after stimulus offset). For units recorded in both species and expression tasks, preferred categories were determined independently.

#### Visualization of decision-variable encoding

Our previous work showed that population-averaged PSTHs in LIP do not monotonically encode the strength of sensory evidence in our face categorization task^10^. Rather, the decision variable is encoded along a curved manifold in the population state space. To compare the temporal dynamics of the decision variable in LIP with TEa, we projected population activity of each area onto an axis aligned with the curved manifold (Fig. 2D, Supplementary Fig. 3B). We used orthogonal canonical correlation analysis (CCA)^72^ to identify a 2D subspace of the population activity that was maximally correlated with two task variables: the decision variable (DV, which is proportional to the signed stimulus strength) and unsigned stimulus strength (stimulus difficulty). As in Okazawa et al. (2021), we focused on a 100-ms window (350-450 ms after stimulus onset) and constructed a population response matrix, **R**, consisting of the trial-averaged, detrended PSTHs across the neural population. The corresponding task variable matrix **P** contained signed and unsigned morph levels. Applying orthogonal CCA on **R** and **P** yielded projection weights **A** and **B**, which map population responses and task variables into a 2D subspace. For any population response vector r⃗, the projection into the subspace is given by:

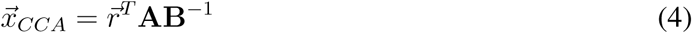

where 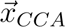 represents the coordinates in the 2D subspace aligned with the DV and stimulus difficulty axes. Fig. 2D illustrates the TEa and LIP activity projected onto the DV axis.

#### Population classifier

To quantify category information in TEa population activity, we performed a population-level classifier analysis. To pool across units recorded in separate sessions, we constructed pseudo-population responses by aligning and combining trials across sessions. To avoid assumptions about specific tuning properties of individual units, we used a template matching approach^51,52^. On each iteration, we randomly selected one trial from each session such that all selected trials shared the stimulus category, stimulus strength (binned as in the PSTH analysis), and the monkey’s choice. The neural activity of these trials was combined to create a pseudo-population response vector for a single test trial. From the remaining trials, we selected correct trials from each category to generate average population activity templates. We then computed the Euclidean distance between the test trial and each template, assigning the test trial to the closer category.

Classification was performed on neural responses in 100-ms sliding windows (1 ms strides), allowing us to track time-varying category information. We repeated this process with randomly sampled test trials, selecting the number of test trials based on the sessions with the fewest trials in the dataset. Classification accuracy was defined as the proportion of trials correctly assigned to their correct category. To estimate the average and standard deviation of this probability, we repeated this analysis 100 times with different random seeds. In Figs. 2C, E, F, and 5C, the solid line represents the average accuracy, and the shaded region indicates the standard deviation across repetitions.

To quantify the duration of ramping in classification performance, we fit a piecewise-linear ramp function to classifier accuracy as a function of time (t):

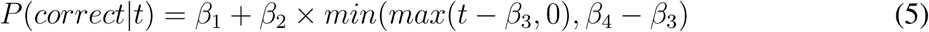

where β_1_ and β_2_ represent baseline accuracy and ramp slope, while β_3_ and β_4_ indicate the onset and plateau times of the ramp. The function was fit separately to each of 1,000 bootstrap samples to estimate variability in ramp timing.

#### Response pulse analysis

To examine how TEa neurons responded to momentary sensory evidence, we exploited the within-trial stimulus fluctuation in our task. We focused on how each stimulus cycle influenced TEa spiking by analyzing residual fluctuations in both stimulus strengths (ΔM*_i_* for each stimulus cycle i; defined as actual morph level in the pulse minus the trial mean stimulus strength) and neural activity (ΔR*_j,t_* for each unit j; defined as firing rate at time t minus the mean firing rate in trials with the same average stimulus strength). We then performed a linear regression to estimate the influence of each pulse on firing rates:

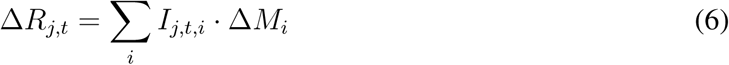

where I*_j,t,i_* denotes the impulse response of unit j at time t to the i-th stimulus pulse. These regression coefficients quantify how strongly each pulse influenced firing at each time point, and the peak weights can be used as a measure of the sensitivity of neural responses to stimulus fluctuations. We performed this analysis on the first four stimulus cycles (i.e., up to the median stimulus duration). The linear regression included a *lasso* regression term, and its coefficient was determined using 5-fold cross-validation. In Fig. 3B, we performed this analysis separately for easy (>15% morph) and difficult (<15% morph) trials to assess how pulse encoding varied over time for different stimulus strengths.

Fig. 3C shows the peak selectivity for each pulse with a finer grouping of trials based on stimulus strength: {0 − 5%, 5 − 15%, 15 − 25%, > 25%}. To estimate the peak amplitude, we fit a Gaussian kernel to the pulse regression weights for the first four stimulus pulses (I*_t,i_* for pulse i = 1..4) as:

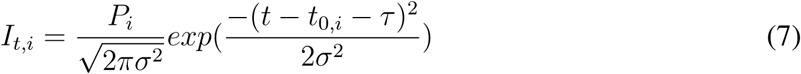

where P*_i_* is the peak amplitude, t_0_*_,i_* is the stimulus onset time of each pulse, and τ and σ are parameters dictating the latency and width of the Gaussian kernel shared across the four pulses. We determined P*_i_*, τ, and σ using a non-linear least squares regression.

### Comparison with encoding models

To better understand what TEa population activity encodes during decision formation, we constructed and compared a series of encoding models built from different subunits that mimic different stages of the decision-making process. These models were evaluated by comparing their outputs to empirical features of the TEa activity: PSTHs (Fig. 4A left) and pulse-triggered responses (Fig. 4A middle and right). Because the drift-diffusion model accurately accounted for the monkey’s behavior (Fig. 1D), we based our simulations on this model, including three processes: representation of sensory inputs, temporal accumulation of evidence, and termination of the accumulation process upon reaching a decision bound.

The simulation began with fluctuating sensory inputs, modeled to match our experimental design (Gaussian fluctuations with 20% morph SD). These inputs were transformed into sensory responses by convolution with a Gaussian temporal kernel (latency: 150 ms; SD: 20 ms), consistent with empirical TEa PSTHs and impulse responses. These computations are captured by the momentary evidence (ME) sub-unit in our models (see below). The output of the ME sub-unit was then integrated over time to produce a decision variable, with accumulation sensitivity k and additive Gaussian noise (σ = 1). This step is captured by the decision-variable (DV) sub-unit in our model. A decision was made when the decision variable reached either an upper (+B) or lower (−B) bound. We selected model parameters (k = 0.5, B = 50) to approximately match the monkey’s psychometric function. The simulation generated two key internal signals: (1) a sensory trace (the convolved input), and (2) a decision variable (the accumulated signal), both of which were used to define the subunits of encoding models described below. Each model simulation consisted of 104 trials per stimulus strength.

#### Construction of encoding models

We constructed several encoding models (Fig. 4C-G) from the ME and DV sub-units (Fig. 4B).

- Base ME sub-unit: Encodes momentary sensory evidence—computed by convolving input fluctuations with a fixed temporal kernel. This sub-unit responds to every stimulus pulse regardless of the DV.
- ME/T sub-unit: Identical to the ME sub-unit, except its responses are set to zero after the DV sub-unit reaches a decision bound, reflecting response termination.
- DV/I sub-unit: Encodes the decision variable by temporally accumulating the ME sub-unit’s outputs. Its activity fluctuates until a bound is reached and remains fixed thereafter. After reaching a decision bound, its response stays at the bound height.
- DV/A sub-unit: It is a variant of the accumulated evidence model, but it encodes the average of sensory evidence obtained up until each moment. We conceived this model because TEa encoded stimulus category more precisely over time (Fig. 2C) without increasing average firing rates (Fig. 2B).
- Hybrid models: A linear combination of the ME and DV sub-units.

In our models, we simplified ME and DV sub-units using symmetric encoding of positive and negative morph levels, which does not fully reflect the asymmetric firing patterns of real neurons (Supplementary Fig. 6). Models that include more complex sub-units or separate sub-units for preferred and non-preferred categories explain the asymmetry of TEa responses. However, these more complex models do not critically change our conclusions.

## Supplementary figures

**Figure S1:**
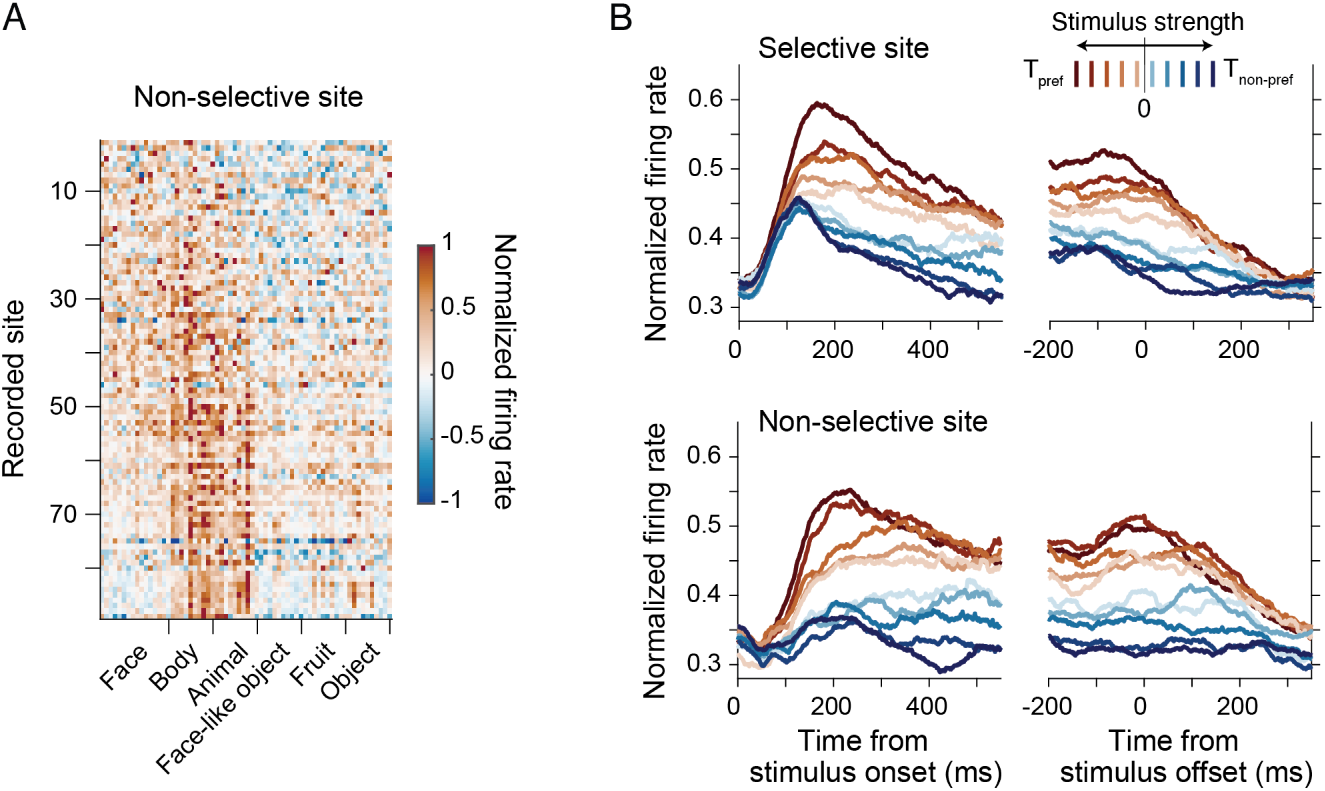
Both face-selective and non-selective sites exhibit similar convergence of responses in the face categorization task. (**A**) Passive visual selectivity for sites not selective to faces. Although these sites did not respond significantly to faces, some showed selectivity for bodies, consistent with their anatomical proximity to face-selective clusters. (**B**) Peri-stimulus time histograms (PSTHs) for face-selective and non-selective sites during the face categorization task. In both populations, responses to the preferred task category (defined by the strongest task-evoked response) gradually converged, indicating similar temporal dynamics.

**Figure S2:**
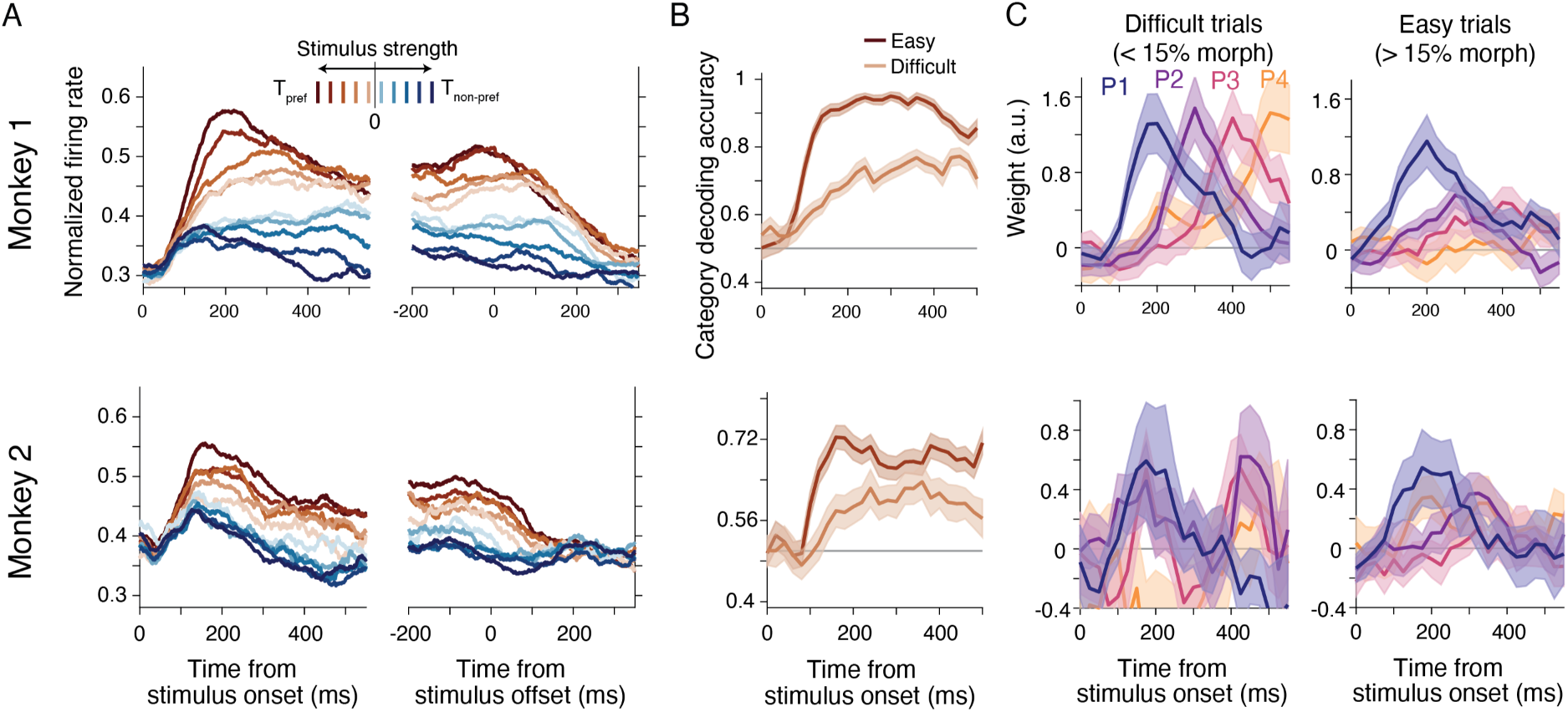
Replication of key results in individual monkeys. **(A-C)** Same analyses as in Fig. 2B, 2C, and 3B, respectively, shown separately for each monkey. All key effects—convergence of firing rates for the preferred category (A), progressive improvement in category decoding (B), and reduction in response to later stimulus pulses on easier trials (C)—were present in both animals. The irregular shape of the results in panel C for monkey 2 arises from suboptimal sampling of stimulus frames.

**Figure S3:**
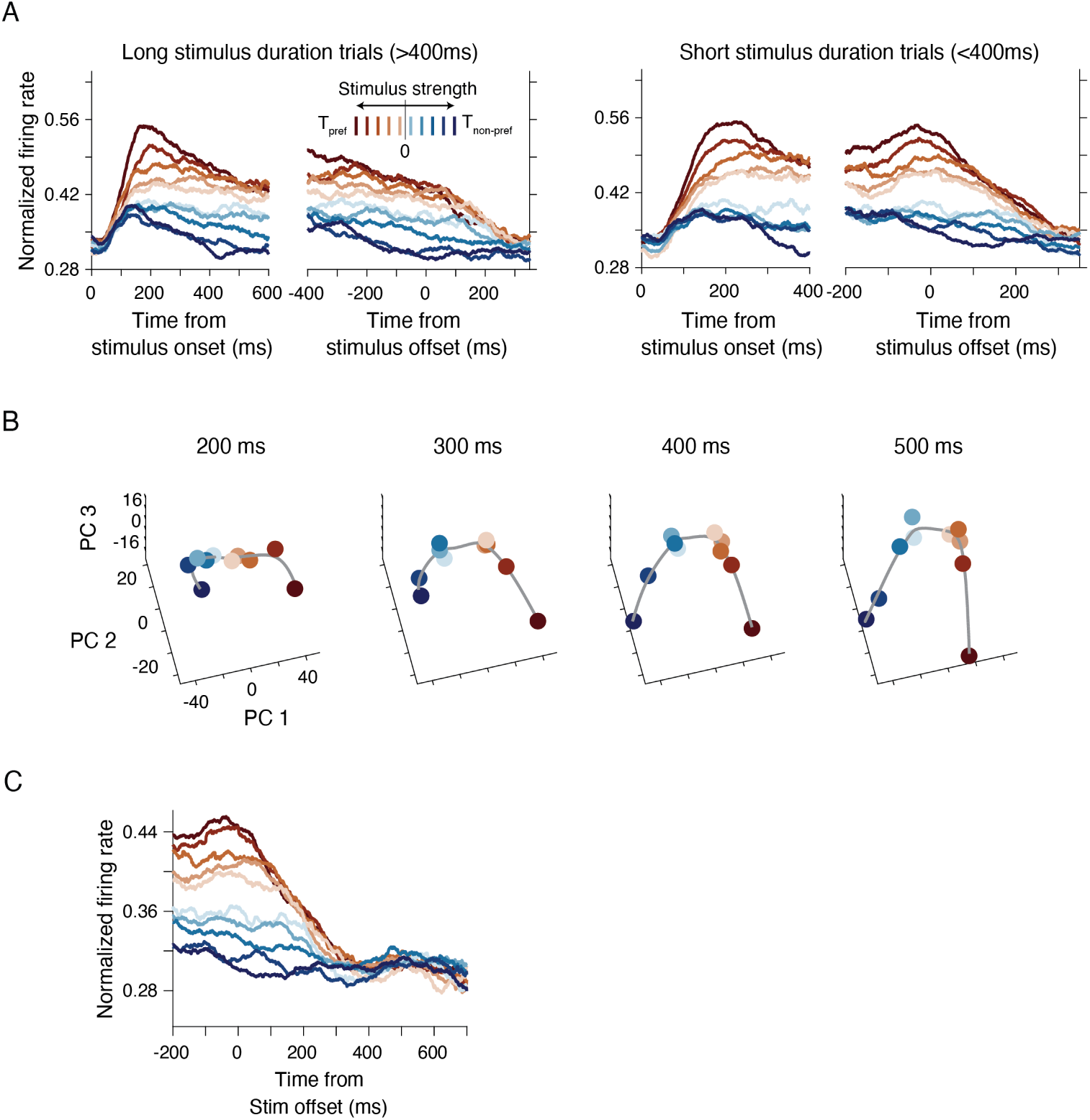
Decision-related signals in TEa emerge during stimulus presentation but are not maintained post-stimulus. (**A**) Convergence of preferred-category PSTHs for trials with short and long stimulus durations. Convergence is more pronounced in long trials, where the extended stimulus duration allows monkeys to reach a decision before stimulus offset. In these trials, convergence begins well before the stimulus ends, indicating that decision-related dynamics in TEa emerge during stimulus presentation, rather than as a post-stimulus effect. (**B**) Principal component analysis (PCA) of TEa population activity. Population activity patterns associated with different morph levels are organized along a curvilinear manifold and become increasingly discriminable during stimulus viewing. This gradual separation mirrors findings in LIP^10^. (**C**) IT population activity does not encode stimulus category or choice during the delay between stimulus offset and behavioral response, suggesting that the DV is not maintained locally in TEa and is likely relayed to other circuits for memory and motor planning.

**Figure S4:**
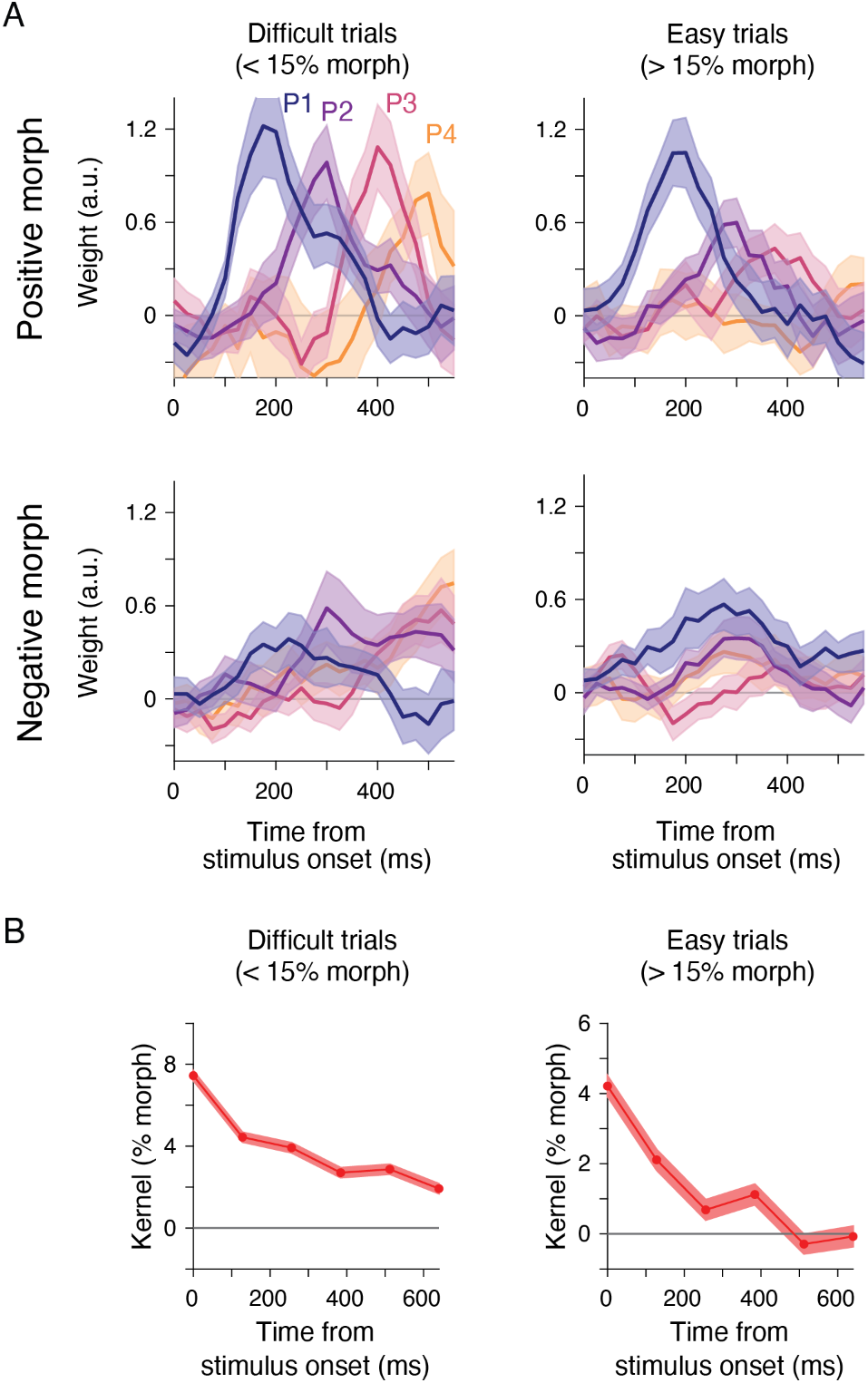
Supporting results for decision termination signals in TEa. (**A**) Reduction in sensitivity to later pulses on easy trials is more prominent when the stimuli belong to the neuron’s preferred category. We separately computed pulse-triggered responses for trials with positive (preferred category) and negative (non-preferred category) morph levels. Conventions are identical to Fig. 3B. (**B**) Behavioral analysis confirmed the reduced influence of late pulses on choice, especially for easy trials, consistent with TEa responses. We performed psychophysical reverse correlation^41,42^ to estimate the temporal weighting of task-relevant sensory evidence. The resulting kernel remained strongly positive throughout stimulus presentation on difficult trials (left), but declined rapidly to near zero after 200 ms on easy trials (right), mirroring the stimulus sensitivity of TEa activity (Fig. 3B). The gradual decline on difficult trials matched quantitative predictions from our extended DDM, in agreement with prior work^10^.

**Figure S5:**
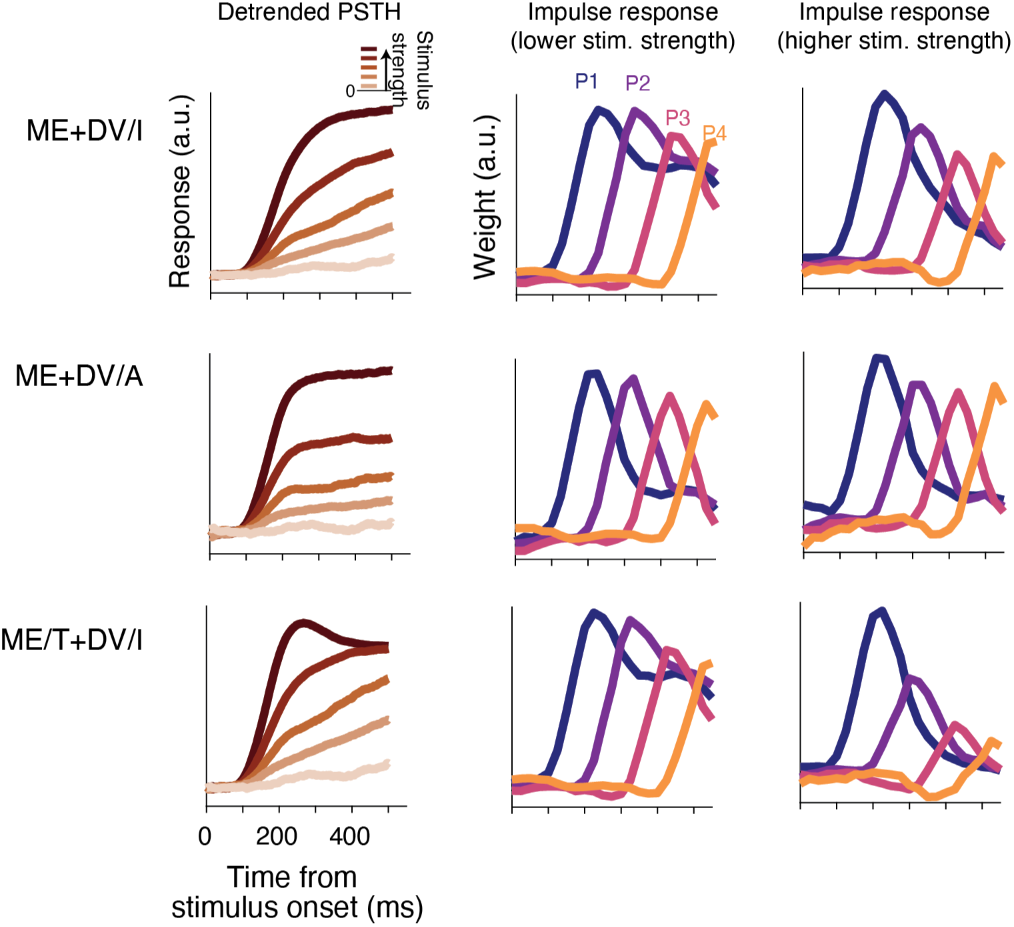
Response dynamics of alternative hybrid sub-unit models. (**A**) ME + DV/I model. (**B**) ME + DV/A model. (**C**) ME/T + DV/I model. Conventions are identical to Fig. 4C-G.

**Figure S6:**
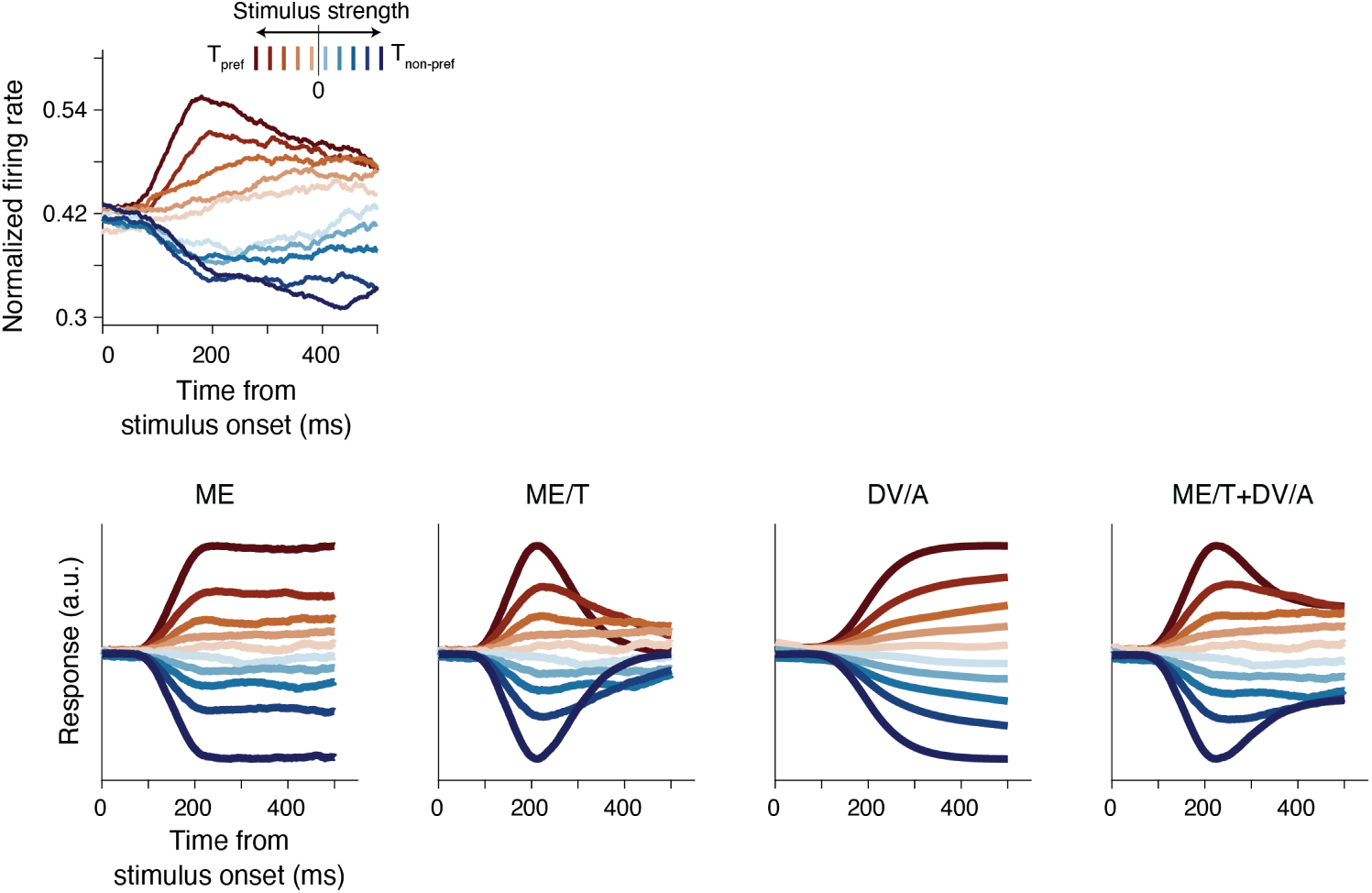
Encoding of non-preferred categories by TEa neurons and simplified sub-unit models. In Fig. 2, we observed an asymmetry in TEa responses to preferred versus non-preferred categories. The simplified sub-unit model in Fig. 4 focused exclusively on responses to the preferred category. Here, we show that the hybrid model with two simple sub-units fails to reproduce the empirical asymmetry in TEa responses. The ME, ME/T, DV/I, and DV/A sub-units all produce symmetrical responses to the two categories. This discrepancy suggests that more complex ME or DV sub-units—or an additional sub-unit—is needed to account for the full range of TEa dynamics.

**Figure S7:**
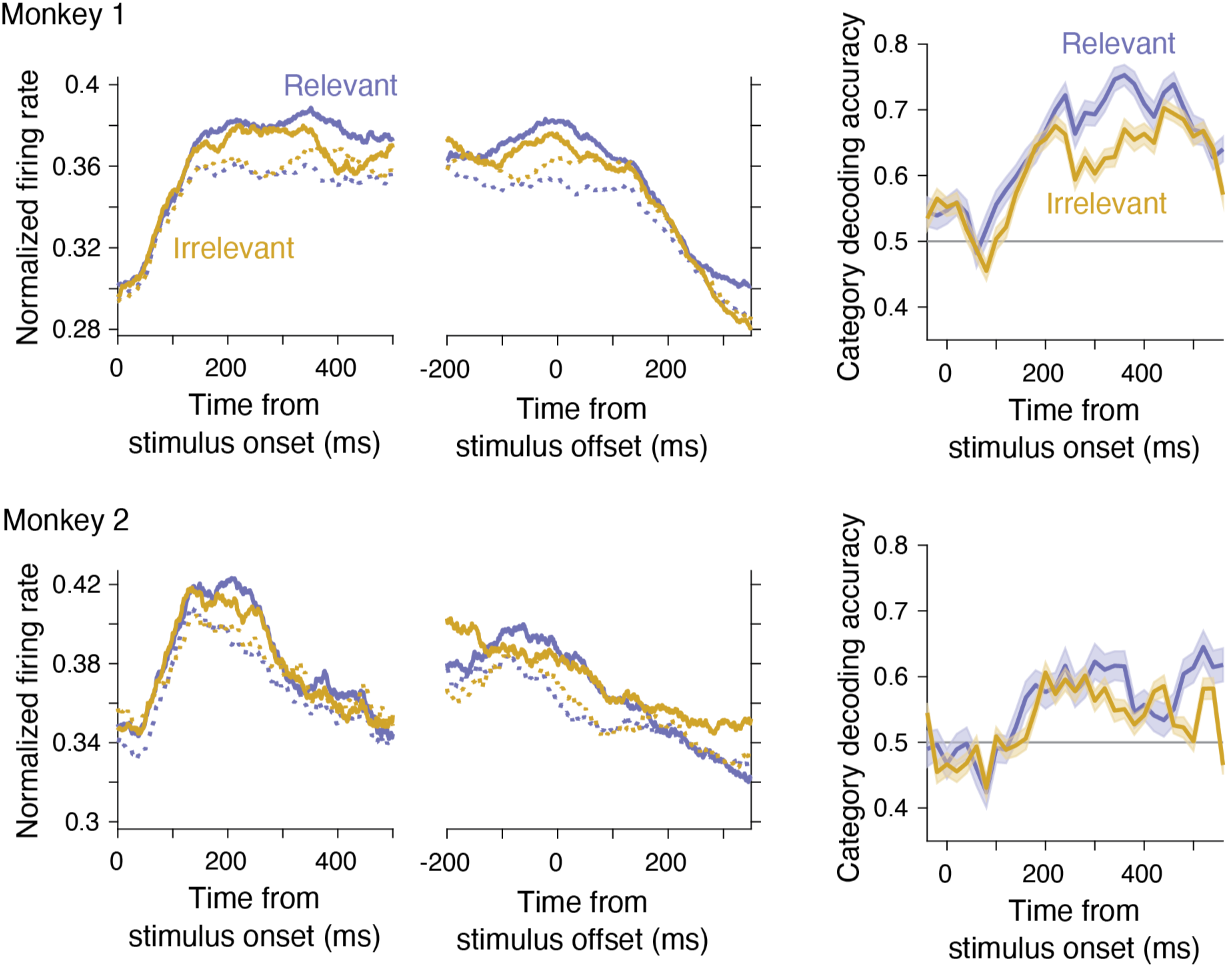
Preferential encoding of task-relevant categories in the TEa of individual monkeys. Conventions are identical to Fig. 5B-C.

## Notes

### Competing Interest Statement

The authors have declared no competing interest.

### Summary of Updates

Abstract, Introduction, Discussion updated for clarification

